# Effect of intracerebroventricular epinephrine microinjection on blood pressure and urinary sodium handling in gestational protein-restricted rat adult male offspring

**DOI:** 10.1101/415778

**Authors:** Bárbara Vaccari Cardoso, Augusto Henrique Custódio, Patrícia Aline Boer, José Antonio Rocha Gontijo

## Abstract

**Background:** In this study, we hypothesized that blunting of the natriuresis response to intracerebroventricularly (i.c.v.) microinjected adrenergic agonists are involved in the development of hypertension in maternal low-protein (LP) intake offspring. **Methods.** Stainless steel cannula was stereotaxically implanted into the right lateral ventricle, by use of techniques reported elsewhere and after that, was evaluated the effect of i.c.v. injection of adrenergic agonists, at increasing concentrations, and of α1 and α2-adrenoceptor antagonists on blood pressure and urinary sodium handling in LP offspring relative to age-matched, normal (NP) protein intake group.

**Results:** We confirmed that epinephrine (Epi) microinjected into the lateral ventricle (LV) of conscious NP rats leads to enhanced natriuresis followed by a reduction in arterial pressure. This response was associated with increased proximal and post-proximal sodium excretion accompanied by an unchanged glomerular filtration rate. The current study showed in both, NP and LP offspring that natriuretic effect of Epi injection into the LV was abolished by prior local microinjection of an α1-adrenoceptor antagonist (prazosin). Conversely, LV α2-adrenoceptor antagonist (yohimbine) administration potentiated the action of epinephrine. The LV yohimbine pretreatment normalized urinary sodium excretion and reduced the blood pressure in LP compared with age-matched NP offspring.

**Conclusion:** These are, as far as we are aware, the first results showing the role of central adrenergic receptors interaction on hypertension pathogenesis in maternal protein-restricted fetal programming offspring. The study also provides good evidence of the existence of central nervous system adrenergic mechanisms consisting of α1 and α2-adrenoceptors, which works reciprocally on the control of renal sodium excretion and blood pressure. Although the precise mechanism of the different natriuretic response of NP and LP rats is still uncertain, these results led us to speculate that inappropriate neural adrenergic pathways might have significant effects on tubule sodium transport, resulting in the inability of the kidneys to control hydrosaline balance, and, consequently, an increase in blood pressure.

## INTRODUCTION

Environmental and genetic factors influence the ontogenetic development, and on adverse circumstance, eventually leading to functional and structural disorders of tissues and organs. In rats, gestational protein restriction is associated with low birthweight, fewer nephrons, and increased risk for the development of heart diseases, kidney dysfunction and metabolic syndrome in adult life [1-7]. We recently demonstrated that maternal low-protein (LP) intake offspring have a lower birth weight, 30% fewer nephrons, and arterial hypertension [3,4,6] when compared with age-matched, normal (NP) protein intake group. Additionally, water and salt balance studies have shown that hypertension in LP offspring is associated with decreased urinary sodium excretion [3–8]. The involvement of the central nervous system (CNS) in the control of blood pressure and, water and salt homeostasis has been demonstrated in several studies [9-12]. It has long been known that there is an association between the CNS and the control of water and salt excretion by the kidneys [1,13,14]. Adrenergic stimulation of the septal area, lateral hypothalamus, subfornical organ, and the anterior region of the third ventricle induces dose-related natriuresis accompanied by, but to a lesser extent, kaliuresis [9,15-21]. On the other hand, studies have shown that electrolytic lesion of the hypothalamic regions in conscious rats reduces salt intake, and the pressor response to cholinergic and noradrenergic microinjection into the median preoptic nucleus [22]. Also, prior findings have been revealed that central α- and β-adrenergic receptors response are involved in CNS hydrosaline homeostasis [9, 22-27].

It has also been postulated that the kidneys are of crucial importance in the pathogenesis of arterial hypertension, as a consequence of primary tubule disorder or renal hemodynamics dysfunction promoting change in tubular sodium and water handling [28-31]. Although the precise mechanism underlying the chronic blood pressure increases in maternal low-protein intake offspring (LP) remains to be elucidated, renal-mediated dysregulation mechanisms of fluid and electrolytes balance are believed to dominate long-term control of arterial blood pressure. Experimental studies support the hypothesis that fetal programming is associated with changes in the tubule sodium transporters in different segments of nephron [32-34]. Research from our laboratory [8] demonstrates that bilateral renal denervation markedly attenuates the increase in arterial pressure and increased tubular sodium excretion in LP offspring. The enhanced urinary sodium excretion in renal denervated LP offspring was accompanied by a significant reduction in proximal tubular sodium reabsorption. This study also demonstrated that impaired dorsal renal ganglia and pelvic neurokinin expression associated with responsiveness of renal sensory receptors are conducive to excess renal reabsorption of sodium and development of hypertension in 16-wk old LP offspring [8]. Given these results, we hypothesized that presumable blunting of the natriuresis response to centrally injected epinephrine α-agonists/antagonists might affect renal tubule sodium transport, resulting in the inability of the kidneys to handle the hydrosaline balance, and consequently, promoting blood pressure enhancement. To test this hypothesis, in this study we evaluated the effect of intracerebroventricular microinjection of adrenergic agonists at increasing concentrations, and α1 and α2 types of adrenoceptor antagonists on blood pressure and urinary sodium handling in LP offspring relative to age-matched normotensive NP counterparts.

## METHODS

### Animals and surgical procedures

The experiments were conducted as described in detail previously [21] on an age-matched female and male rats of sibling-mated Wistar *HanUnib* rats (250–300 g) that were allowed free access to water and standard rodent chow (Nuvital, Curitiba, PR, Brazil). The Institutional Ethics Committee (CEUA/UNICAMP #2766/01) approved the experimental protocol, and the general guidelines established by the Brazilian College of Animal Experimentation were followed throughout the investigation. The environment and housing presented the right conditions for managing their health and well-being during the experimental procedure. Immediately after weaning at three weeks of age, animals were maintained under controlled temperature (25°C) and lighting conditions (07:00–19:00 h) with free access to tap water and standard laboratory rodent chow (Purina rat chow: Na^+^ content: 135 ± 3μEq/g; K^+^ content: 293 ± 5μEq/g), for ten weeks before breeding. It was designated day 1 of pregnancy as the day in which the vaginal smear exhibited sperm. The dams were maintained on isocaloric rodent laboratory chow with standard protein content [NP], (17% protein) or low protein content [LP] (6% protein) diet, *ad libitum* intake, throughout the entire pregnancy. The NP and LP maternal food consumption were determined daily (subsequently normalized for body weight), and body mass of dams was recorded weekly in both groups. All maternal groups returned to the standard laboratory rodent chow after delivery. The male pups weaned for three weeks, and only one offspring from each litter was used for each experiment. The male offspring also were followed and maintained with standard rodent chow under a controlled temperature (25°C), a 12 h light–dark cycle (07:00–19:00), and housed in cages with four rats until 16 weeks of age. The NP and LP offspring were handled weekly by a familiar person and followed until 16 weeks of age when physiological tests were performed. The litter number was determined, the anogenital distance was measured and, male offspring weighed at birth and weekly after that. Male NP and LP offspring (from 180 to 250 g) were chronically instrumented with a cerebral right lateral ventricle (LV) guide cannula and kept under controlled temperature (25°C) and lighting conditions (07:00 h-19:00 h) in individual cages. Briefly, the animals were anesthetized with a mixture of ketamine (75 mg.kg-1 body mass, and xylazine 10 mg.kg-1 body mass), injected intraperitoneally (i.p.). Once the corneal and pedal reflexes were absent, a stainless steel cannula was stereotaxically implanted into the right LV, by use of techniques reported elsewhere and pre-established coordinates: anteroposterior +0.2 mm from bregma; lateral +1.5 mm; and vertical −4.5 mm [10,21,26,27]. After stereotaxic surgery, rats were allowed one-week recovery before testing for cannula patency and position. The position of the cannula was visually confirmed by infusion of blue Evans 2 % through the LV cannula at the end of the experiment (Figure 1). Data from animals with incorrectly placed cannulas were excluded from statistical analysis.

**Figure 1.**
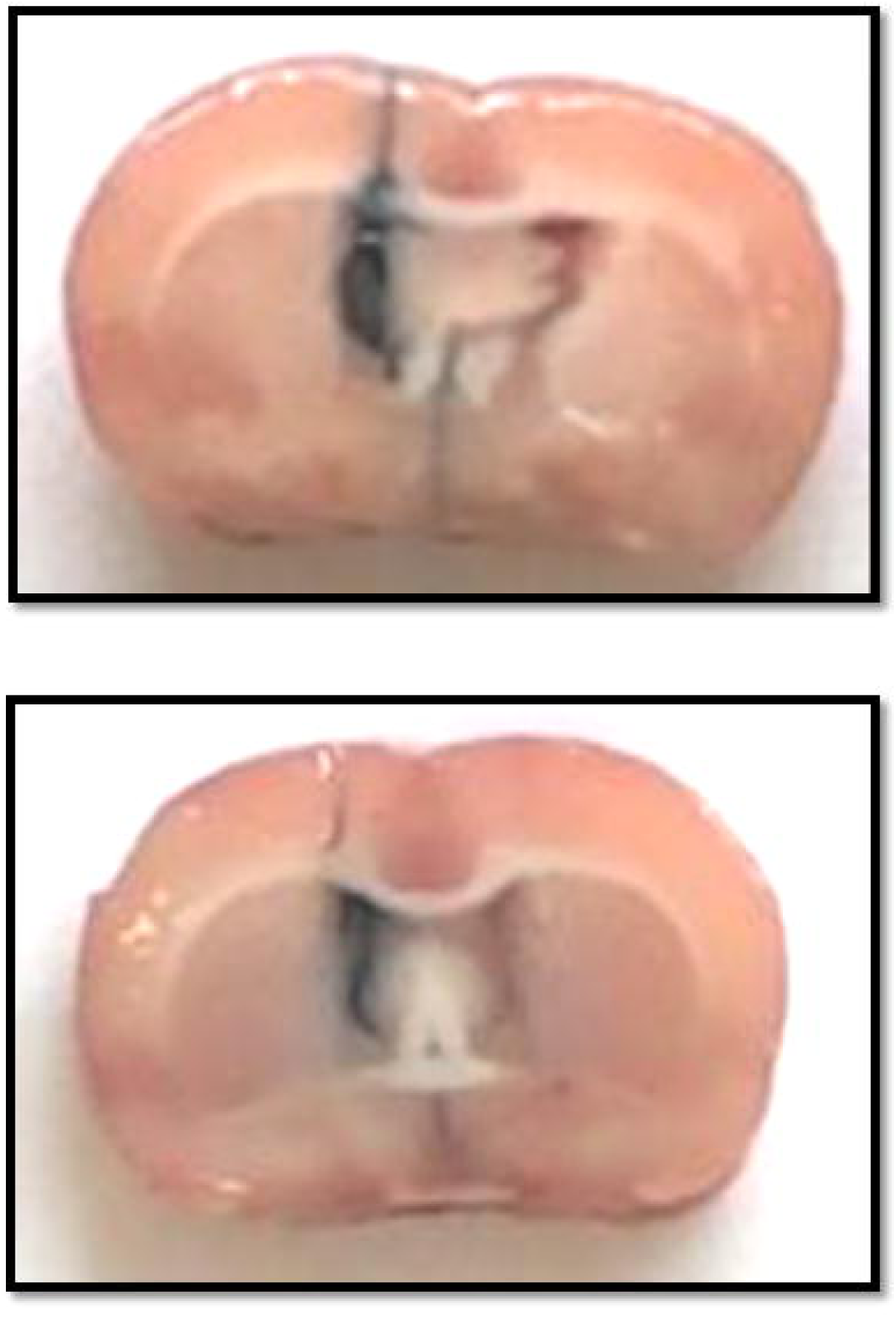
Histological verification of the cerebral right lateral ventricle (LV) guide cannula.

### Renal function test

The effect of intracerebroventricular adrenergic (epinephrine, Epi) microinjections on glomerular filtration and tubular sodium handling was assessed by creatinine and lithium clearance at 16 weeks of age (NP, n = 10 and LP, n = 10), in unanesthetized, unrestrained male offspring. Briefly, fourteen hours before the renal test, 60μmol LiCl/100 g^−1^ body weight was given by gavage. At 8:00 a.m., after an overnight fast with free access to water, each animal received tap water by gavage (5 % of body mass), followed by the same volume 1-h later. Immediately before experiments, the animals’ (NP and LP) water supply was removed from the home cage. The indwelling obturator was replaced by a 30-gauge stainless steel injector at the end of PE-10 tubing connected to a 10μl Hamilton syringe wholly filled with the test solution. Afterward, 30 min after the second volume of water, epinephrine (Epi, *n* = 9 to 10) for each dose, Sigma Chemical Co., MO, USA), or a similar amount of 0.15 M NaCl (vehicle, *n* = 8) was microinjected into the LV in a volume of 3μl, at different concentrations of epinephrine (Epi: 0.1, 0.3 and 1.0µmol) diluted in saline, of one NP or LP offspring from each litter. The drug was administered by a 10-μl Hamilton micro-syringe, and urine spontaneously voided over five 30 min-period, one before (−30min) and four after i.c.v. drug administration (30 to 120 min), was collected into graduated centrifuge tubes and measured gravimetrically. To examine the effect of α-adrenergic receptor antagonist after LV administration of 0.3µmol Epi, rats were randomly assigned to one of specific experimental group and a selective α1-adrenoceptor antagonist (4-nmol prazosin, Pz; n=9, Sigma Chemical Co., MO, USA), α2-adrenoceptor antagonist (4-nmol Yohimbine, Yo; n=10, Sigma Chemical Co., MO, USA) or vehicle (0.15M NaCl, n=8) was LV microinjected the 30-min before the agonist in a volume of 1μl. At the end of the experiment, blood samples were drawn by cardiac puncture from anesthetized rats and urine, and plasma samples were collected for analysis [26,27,29,35].

### Blood pressure measurement

The systolic arterial pressure was measured in conscious and previously trained offspring at 8 weeks to 16 weeks of age in NP (n = 10 to 36) and LP (n = 10 to 32) offspring. In a subgroup a part of animals, the systolic blood pressure (SBP) was measured 30-min before and after LV administration of 0.3µmol Epi and/or receptor antagonists administration followed for two subsequent periods of 60 minutes, in conscious NP and LP offspring, employing an indirect tail-cuff method using an electrosphygmomanometer combined with a pneumatic pulse transducer and amplifier (IITC Life Science BpMonWin Monitor Version 1.33). This indirect approach enabled repeated measurements with close correlation (correlation coefficient = 0.975), compared with direct intra-arterial recording [3,4,8,29,35]. The mean of three consecutive readings represented the blood pressure.

### Data presentation and statistical analysis

All numerical results are expressed as the mean ± standard deviation (SD) or median and quartile deviation as appropriate. Plasma and urine sodium, potassium, and lithium concentrations were measured by flame photometry (Micronal B262, São Paulo, Brazil). The integrated renal fractional sodium and potassium excretions after 0.3µmol Epi LV microinjection, i.e., the total area under the curve (AUC, %.120 min^-1^) was calculated by the trapezoidal method. Plasma and urine sodium, potassium and lithium concentrations were measured by flame photometry (Micronal, B262, São Paulo, Brazil), while the creatinine concentrations and plasma osmolality were determined spectrophotometrically (Instruments Laboratory, Genesys V, USA) and by Wide-range Osmometer (Advanced Inst. Inc, MA, USA), respectively. After that, the animals were anesthetized with ketamine and xylazine injected intraperitoneally and sacrificed by cardiac puncture; urine and plasma samples were stored for analysis. Creatinine clearance (*C*_*cr*_) was assessed to estimate glomerular filtration rate, and lithium clearance (_*CLi*_) was used to determine proximal tubule output [3,4,8,29,35]. Fractional sodium excretion (*FE*_*Na*_) was calculated as *C*_*Na*_/*C*Cr × 100, where *C*_Na_ is sodium clearance, and *C*Cr is creatinine clearance. Fractional proximal (*FEP*_*Na*_) and post-proximal (*FEPP*_Na_) sodium excretion were calculated as *C*_Li_/*C*_Cr_ × 100 and *C*_*Na*_/*C*_Li_ × 100, respectively. Data obtained over time were analyzed by the use of repeated measures two-way ANOVA. *Post-hoc* comparisons between selected means were performed with Bonferroni’s contrast test when initial ANOVA indicated statistical differences between experimental groups. When relevant, comparisons involving only two means within or between groups were achieved by use of a Student’s *t*-test. Statistical analysis was performed with GraphPad Prism 5.01 for Windows (1992-2007 GraphPad Software, Inc., La Jolla, CA, USA). The level of significance was set at *P*<0.05. Availability of data and material: Available in Unicamp Repository at http://repositorio.unicamp.br/bitstream/REPOSIP/312742/1/Cardoso_BarbaraVaccari_M.pdf and, https://sucupira.capes.gov.br/sucupira/public/consultas/coleta/trabalhoConclusao/viewTrabalhoConclusao.jsf?popup=true&id_trabalho=1479247

## RESULTS

### Dams and offspring parameters

Table 1 shows serum sodium, lithium, and potassium levels from NP and LP offspring, with no significant differences in NP rats compared with the LP group. In general, water and food sodium intake and, plasma osmolality were similar in male offspring of NP and LP groups when normalized by body weight (Table 1). Gestational protein restriction did not significantly change the pregnant dams’ body mass during gestation (Figure 2A). Also, it did not affect the number of offspring per litter and the proportion of male and female offspring (p=0.3245). The birthweight of LP male pups (n=26) was significantly reduced compared with NP pups (n=27) (6.238 ± 0.2288 g vs. 7.200 ± 0.05163 g; *P* = 0.0001), respectively (Figure 2B). The body mass of 12 days weaning LP pups (21 ± 0.5 g, n=18) remained lower than compared age-matched NP pups (24 ± 0.6 g, n=19) (*P* = 0.0017) (Table 1). The body mass of LP pups (67.9 ± 0.99 g, n = 17) remained lower than age-matched NP pups (72.10 ± 1.42 g, n = 20) until weaning at 21 days after birth (*P* = 0.05) (Figure 1C). However, between 4 weeks and 16 weeks of age, the body weight of LP and NP rats was not significantly different (*P* = 0.5170) (Figure 2C).

**Table 1.** Serum sodium, lithium, and potassium levels and sodium and water intake and plasma osmolality in maternal normal protein intake (NP) offspring compared with maternal low-protein intake (LP) offspring (n = 10 animals for each group). The data represent the means ± SEM. The level of significance was set at *P ≤ 0.05 (one-way ANOVA or Student’s t-test).

| *Groups/Parameters* | *Na<sup>+</sup> (mM)* | *Li<sup>+</sup> (μM)* | *K<sup>+</sup> (mM)* | *Na<sup>+</sup>Intake*<br>*(mmol.wk<sup>-1</sup>,100g<sup>-1</sup> bw)* | *H<sub>2</sub>O Intake*<br>*(ml.100g<sup>-1</sup>bw)* | *Plasma Osmolality*<br>*(mOsm.kg<sup>-1</sup>H<sub>2</sub>O)* |
| --- | --- | --- | --- | --- | --- | --- |
| *NP (n=10)* | 139 ± 4.6 | 86 ± 19 | 4.5 ±0.7 | 14.3 ± 2.3 | 23.6 ± 8.2 | 295 ± 8.2 |
| *LP (n=10)* | 141 ± 4.3 | 81 ± 18 | 4.3± 0.5 | 13.5 ± 2.5 | 23.7 ± 4.8 | 297 ± 7.7 |

**Figure 2.**
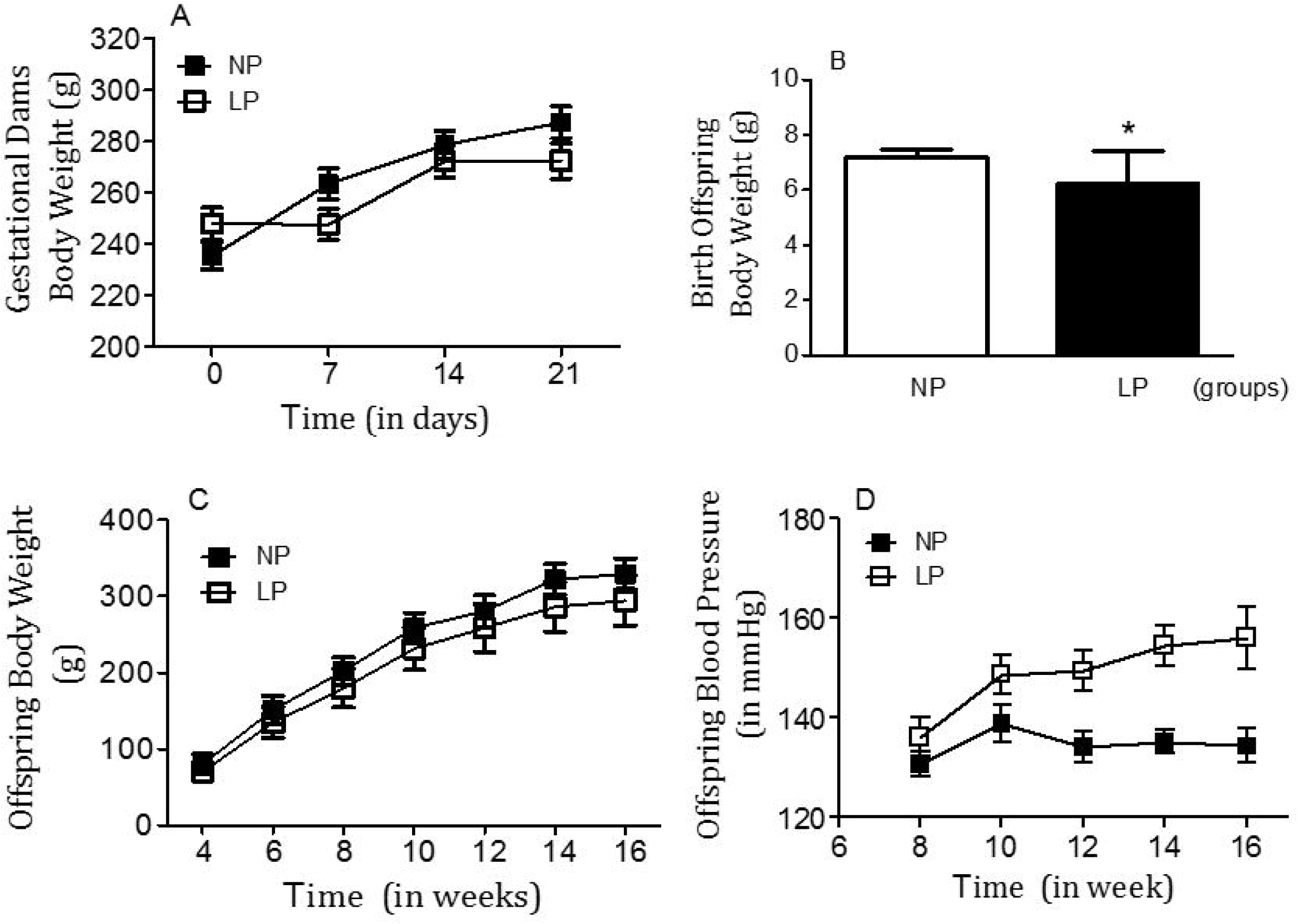
Gestational dams’ body weights (A), offspring body weight at birth (B) and (C) from 4-day to 16-week-old (in grams) and offspring blood pressure from 8 to 16 week-old (in mmHg) NP compared to age-matched LP offspring. The results are expressed as means ± SD. Data were analyzed using a two-way ANOVA test with *post-hoc* comparisons by Bonferroni’s contrast test. The level of significance was set at *P < 0.05.

### Blood pressure measurement

Blood pressure and renal function data (expressed as mean ± SD) for 16-week-old NP (n = 10 to 32 offspring for each control group) and LP (n = 10 to 36 for each experimental group) offspring are summarized in Figures 3 to 7 and Table 1. As shown in Figure 2D, tail systolic arterial pressure (in mmHg) was significantly higher in LP offspring compared with NP offspring between 8 weeks and 16 weeks of age (*P* = 0.0001). The changes in systolic blood pressure from 8 weeks to 16 weeks of age were as follows: 8 weeks: LP, 136.2 ± 4.04 mmHg vs. NP, 130.7 ± 2.43 mmHg, *P* = 0.0001; 16 weeks: LP, 156.1 ± 6.25 mmHg vs. NP, 134.4 ± 4.04 mmHg, *P* = 0.0001 (Figure 2D). This study also revealed in NP offspring a rapid, transient, but considerable blood pressure decrease after LV microinjection of 0.3µmol Epi (Figure 4A and 4B; NP, n=9; *P* = 0.0001) an effect which, in turn, was significantly attenuated by α1-adrenoceptor antagonists (4 nmol prazosin, n=10; *P* = 0.2596) (Fig. 4A) and, was unchanged by the α2-receptor antagonist (NP: 4 nmol yohimbine, n=10; *P* = 0.1395) (Fig. 4B). No blood pressure changes, to LV microinjection of 0.3µmol Epi or Epi+α1-adrenoceptor antagonists (*P* = 0.0629 and *P* = 0.3051, respectively), was observed in 16-week old LP offspring. However, LV microinjection of 0.3µmol Epi+α2-adrenoceptor antagonists causes a significant reduction in blood pressure in LP offspring (*P* = 0.0007) (Fig. 4B).

**Figure 3.**
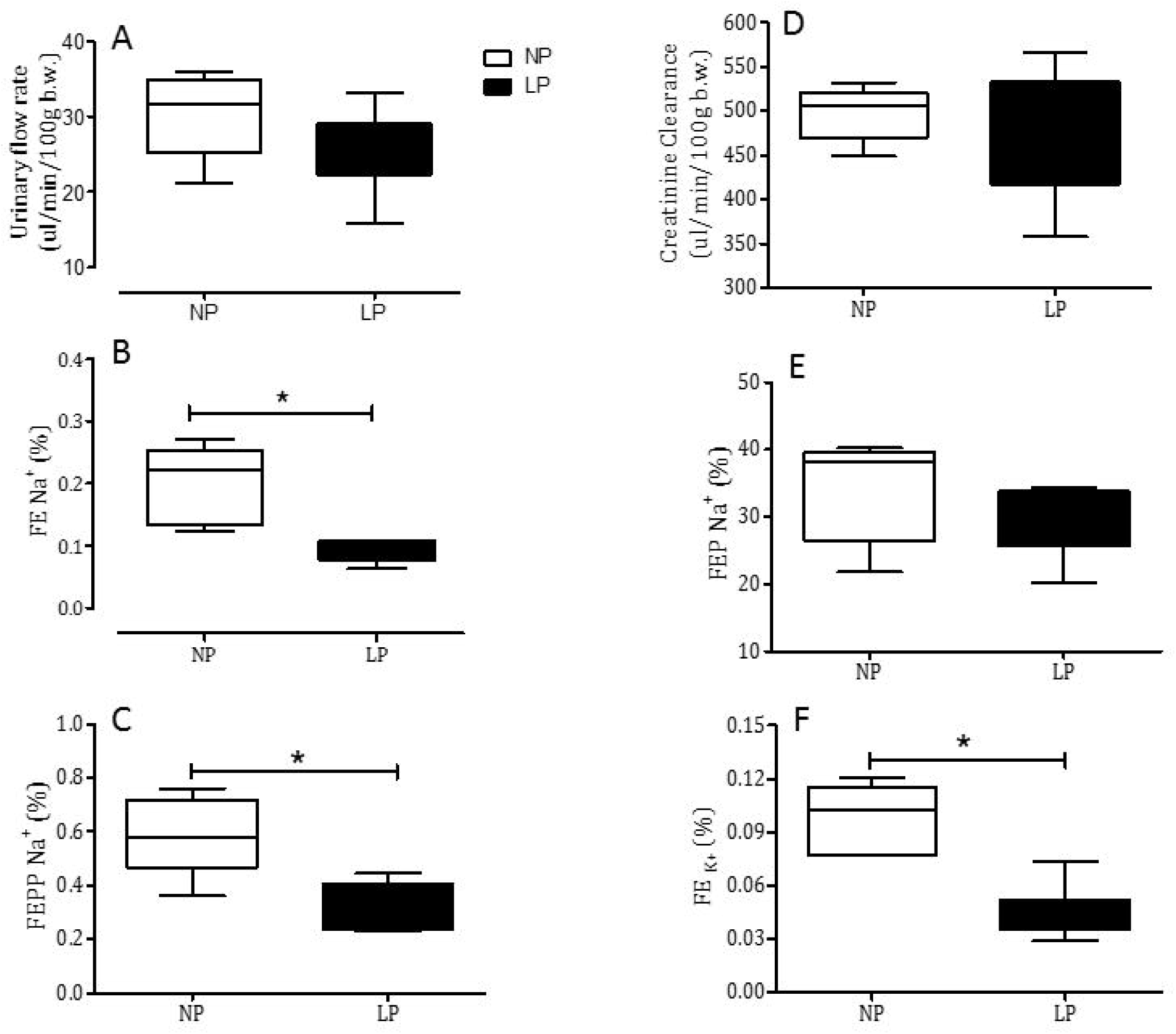
Renal function studies using urinary flow rate (Panel A), fractional sodium excretion (FENa+, panel B), post-proximal (FEPPNa+, panel C), proximal (FEPNa+, panel E), fractional sodium excretion, creatinine clearance (CCr, panel D), and fractional potassium excretion (FEK+, panel F), in male 16-week-old NP compared to age-matched LP (n = 10 for each group). Results are expressed as median and quartile deviation. Data were analyzed using a one-way ANOVA test. The level of significance was set at *P < 0.05.

**Figure 4.**
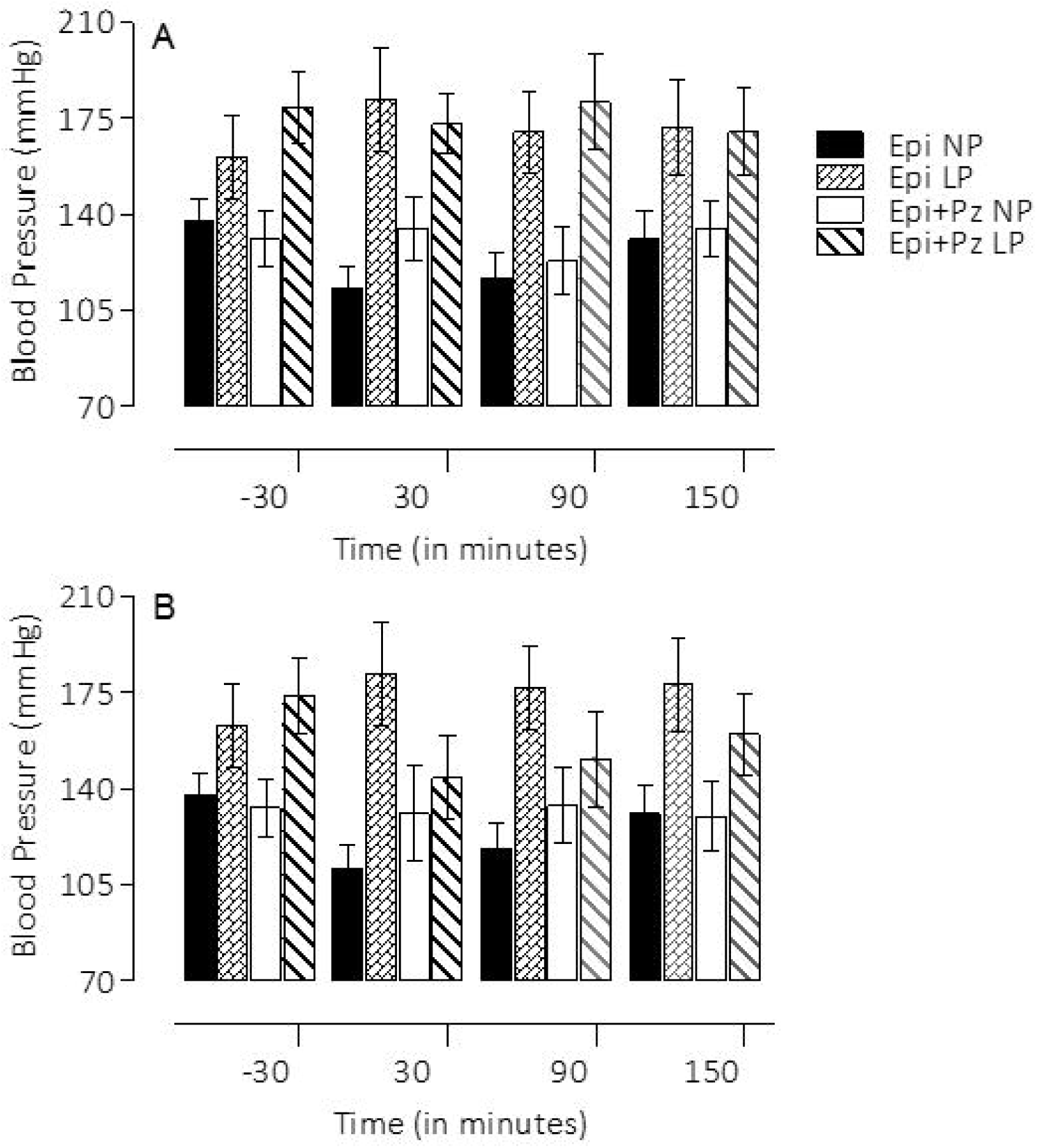
Effect of the lateral ventricle (LV) microinjection of agonist 0.3-µmol epinephrine, and their antagonists, 4-nmol prazosin (A) and 4 nmol yohimbine (B), on blood pressure among gestational normal (NP) protein intake group relative to maternal protein-restricted offspring (LP). Data were analyzed using a two-way ANOVA test with *post-hoc* comparisons by Bonferroni’s contrast test. The level of significance was set at *P < 0.05.

### Renal function data - Dose-response curve for Epi-induced sodium excretion response

Renal function in 16-week-old NP and LP offspring is summarized in Figure 2. No changes in plasma sodium, lithium, or potassium levels were observed in any of the experimental groups (Table 1). Also, the urinary flow rates (*P* = 0.0782) and the glomerular filtration rate, (*P* = 0.1959), estimated by *CCr*, did not significantly differ between the NP and LP offspring (respectively, Figure 4A and 4D). Fractional urinary sodium excretion (FENa, Figure 4B) in 16-week old LP rats was significantly reduced in LP offspring relative to age-matched NP rats (16-week-old LP: 0.089 ± 0.006% vs. NP: 0.199 ± 0.028%; *P* = 0.006). The decreased FENa in LP rats was accompanied by a significant reduction in FEPPNa (16-week-old LP: 0.324 ± 0.031% vs. NP: 0.589 ± 0.07%; *P* = 0.0013) and FEK (16-week-old LP: 0.046 ± 0.006% vs. NP: 0.0973 ± 0.008%; *P* = 0.0002), but not by FEPNa when compared with age-matched NP control rats (Figures 4C, 4E and 4F). Lateral ventricular (LV) microinjections, in a dose-dependent fashion, of 0.1, 0.3 and 1.0µmol Epi diluted in 3μl volume promoted an increase in urinary sodium and potassium excretion over 120 min in 16-week-old NP; this effect was significantly (*P* = 0.0001) attenuated in age-matched LP offspring (Fig. 5 and 6). After dose-response experiments, a dose of 0.3µmol Epi was selected as adequate for the rest of the study and the results showed as the total area under the curve (AUC, %.120 min-1). The effect of LV 0.3µmol Epi on increasing renal fractional sodium excretion was significantly higher for NP than in LP offspring (*P* ≤ 0.001) (Fig. 6). As depicted in the Figures 6 and 7, a consistent increase of FENa among NP offspring was accompanied by significant enhancement of proximal (from basal 38.3 ± 9.5 to 73.3 ± 21.2 %, *P* = 0.0002) and post-proximal (from basal 38.3 ± 9.5 to 73.3 ± 21.2 %, P = 0.0245) sodium excretion; for LP group, a smaller but significant increase in proximal (from 42.9 ± 8.3 to 54.7 ± 7.6 %) but not in post-proximal (from 2.36 ± 0.38 to 5.12 ± 0.62 %) sodium excretion was observed (*P* = 0.002 and *P* = 0.001, respectively) (Fig. 5). The increase occurred in association with unchanged *CCr*. The effects of LV antagonist adrenoceptors administration on urinary sodium excretion was also studied for both offspring groups, bearing implanted cannulas. This study revealed the participation of LV α1 and α2 - adrenergic receptors in the regulation of renal sodium and potassium excretion. The increased natriuresis and kaliuresis response of 0.3µmol Epi microinjection into LV in 16-week-old NP rats but less, in age-matched LP offspring, was significantly attenuated by previous local injection of an α1-adrenergic antagonist (4 nmol prazosin) (*P* = 0.001) (Fig. 7). Conversely, to observed in NP rats, the current findings also support the observation that LV pre-injection of 4-nmol yohimbine, an α2-adrenergic antagonist, synergically potentiated the action of 0.3µmol Epi LV administration (Fig. 8), on renal sodium excretion in LP offspring. Note that prazosin (Fig. 7) significantly inhibited fractional sodium excretion whereas yohimbine (Fig. 8) enhanced fractional sodium excretion. Surprisingly, the LV yohimbine pretreatment normalized urinary sodium excretion by LP compared with age-matched NP offspring at 16 weeks of age. (*P* ≤ 0.001, Fig. 8).

**Figure 5.**
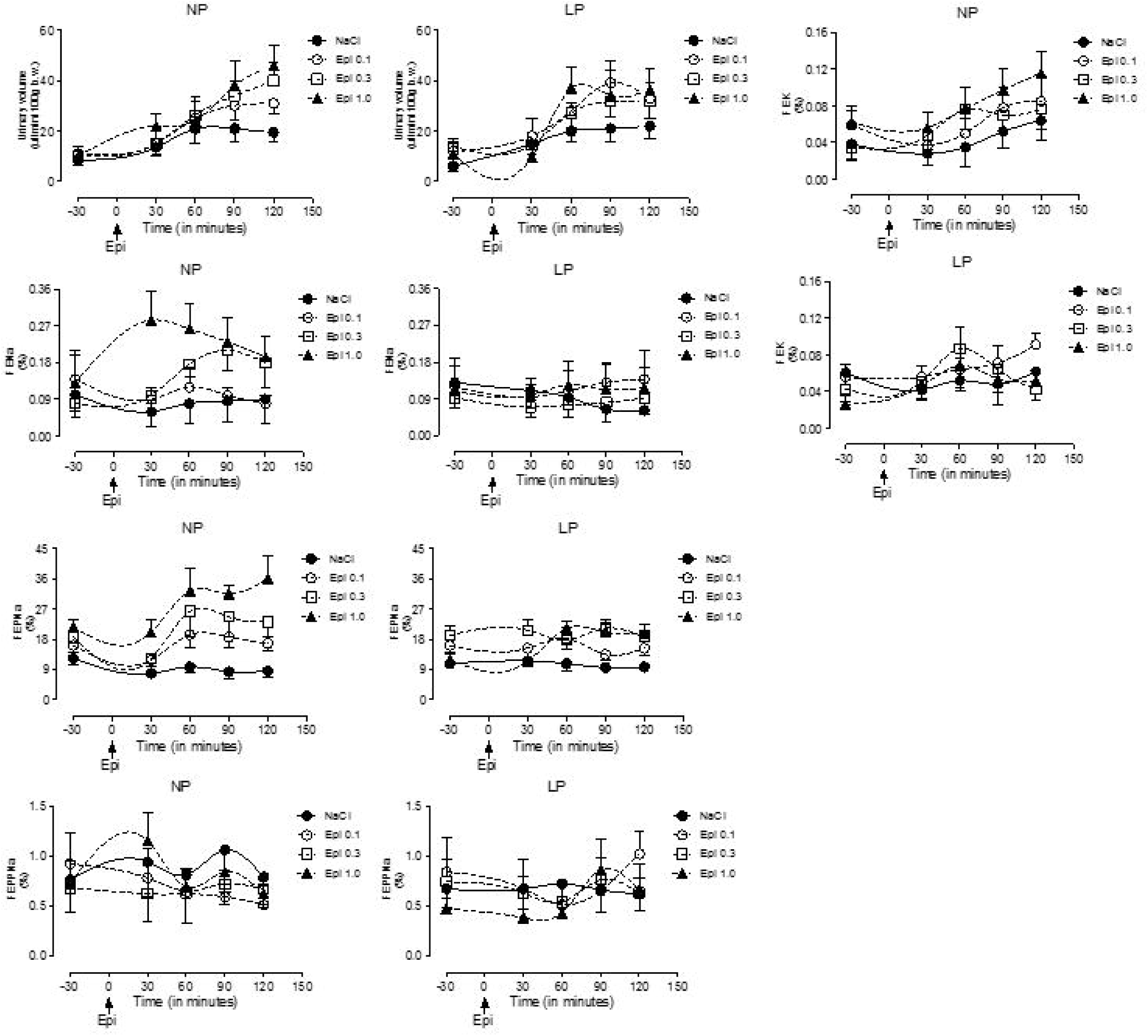
Dose-response curve to LV microinjection of agonists at different concentrations of epinephrine (Epi: 0.1, 0.3 and 1.0µmol) in a volume of 3μl, on urine volume, fractional sodium excretion (FENa), proximal (FEPNa), post-proximal (FEPPNa) fractional sodium excretion, and fractional potassium excretion (FEK), in NP compared to age-matched LP offspring. Results are reported as mean ± SD. Data were analyzed using a two-way ANOVA test with post-hoc comparisons by Bonferroni’s contrast test. The level of significance was set at *P < 0.05. The study was controlled by the lateral ventricle (LV) microinjection of 0.15 M NaCl.

**Figure 6.**
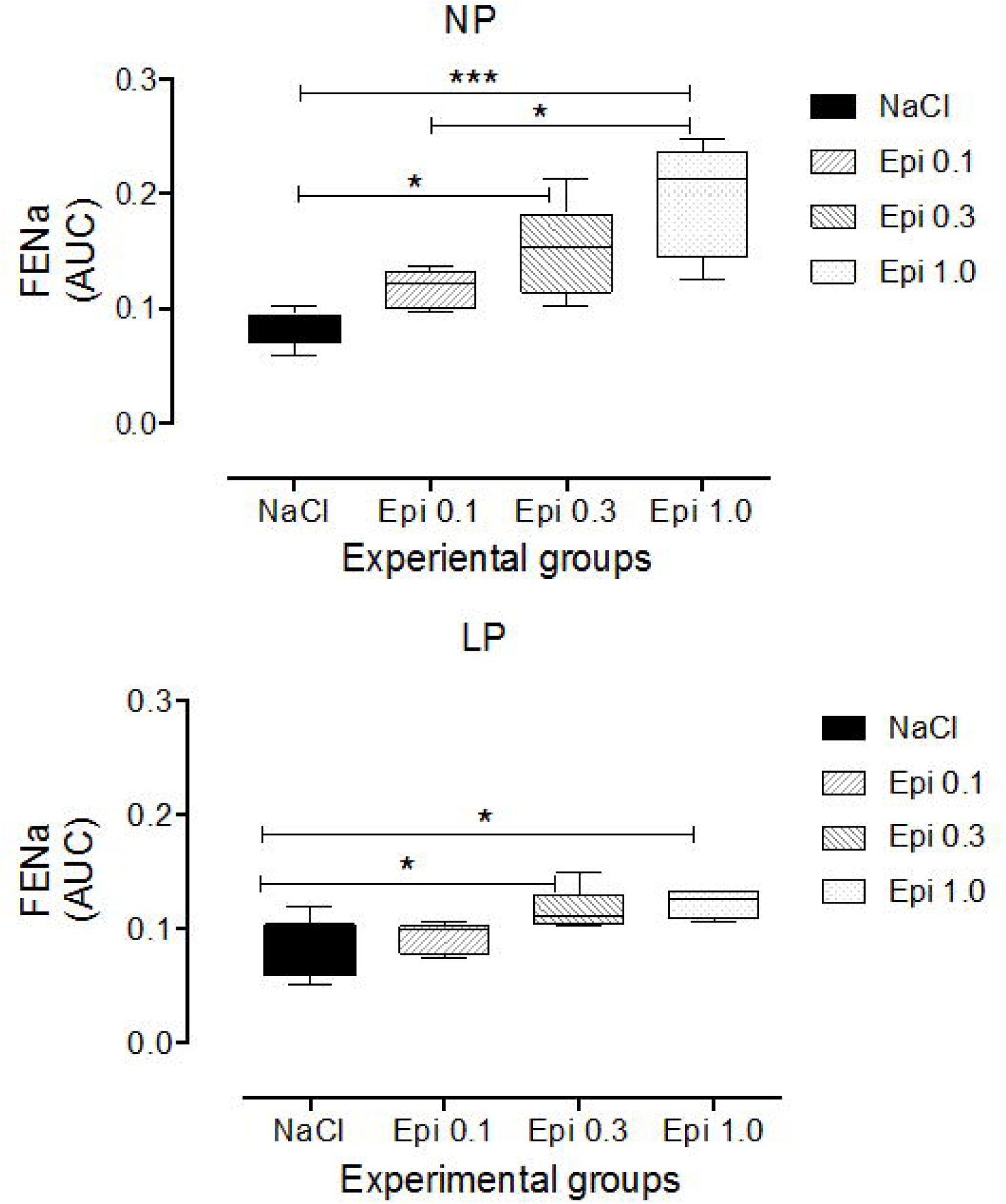
Dose-response curve to LV microinjection of agonists at different concentrations of epinephrine (Epi: 0.1, 0.3 and 1.0µmol) in a volume of 3μl, on fractional sodium excretion (FENa), expressed as mean ± SD of the total area under the curve (AUC, %.120 min-1). Data were analyzed using a two-way ANOVA test with post-hoc comparisons by Bonferroni’s contrast test. The level of significance was set at *P < 0.05. The study was controlled by the lateral ventricle (LV) microinjection of 0.15 M NaCl.

**Figure 7.**
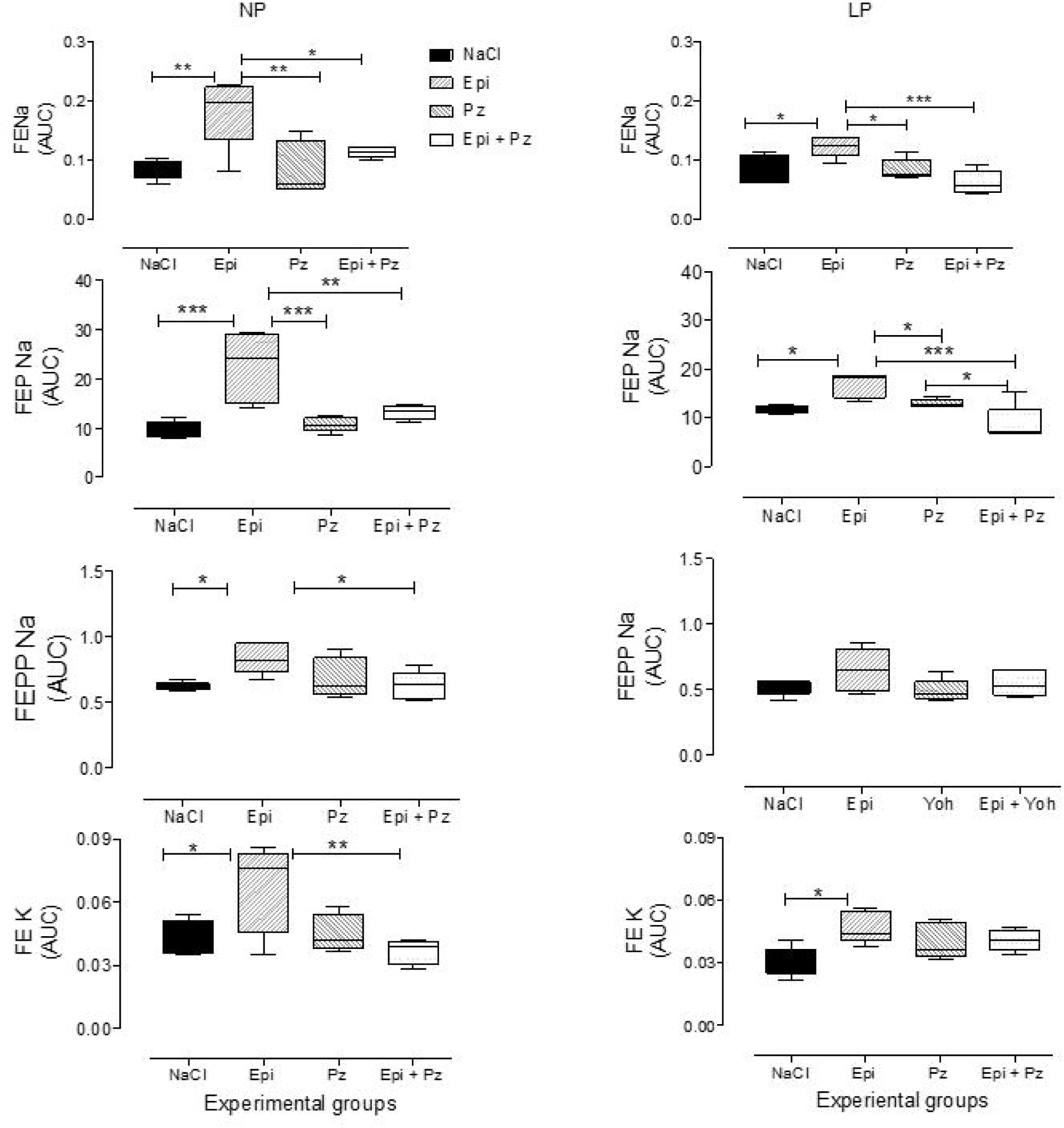
Effect of the lateral ventricle (LV) microinjection of agonist 0.3-µmol epinephrine and/or their α1-adrenoceptor antagonist, 4-nmol prazosin (Pz), on fractional sodium excretion (FENa), proximal (FEPNa), post-proximal (FEPPNa) fractional sodium excretion, and fractional potassium excretion (FEK), in NP, compared to age-matched LP offspring. Results are reported as mean ± SD. Data were analyzed using a two-way ANOVA test with post-hoc comparisons by Bonferroni’s contrast test. The level of significance was set at *P < 0.05. The study was controlled by the lateral ventricle (LV) microinjection of 0.15 M NaCl.

**Figure 8.**
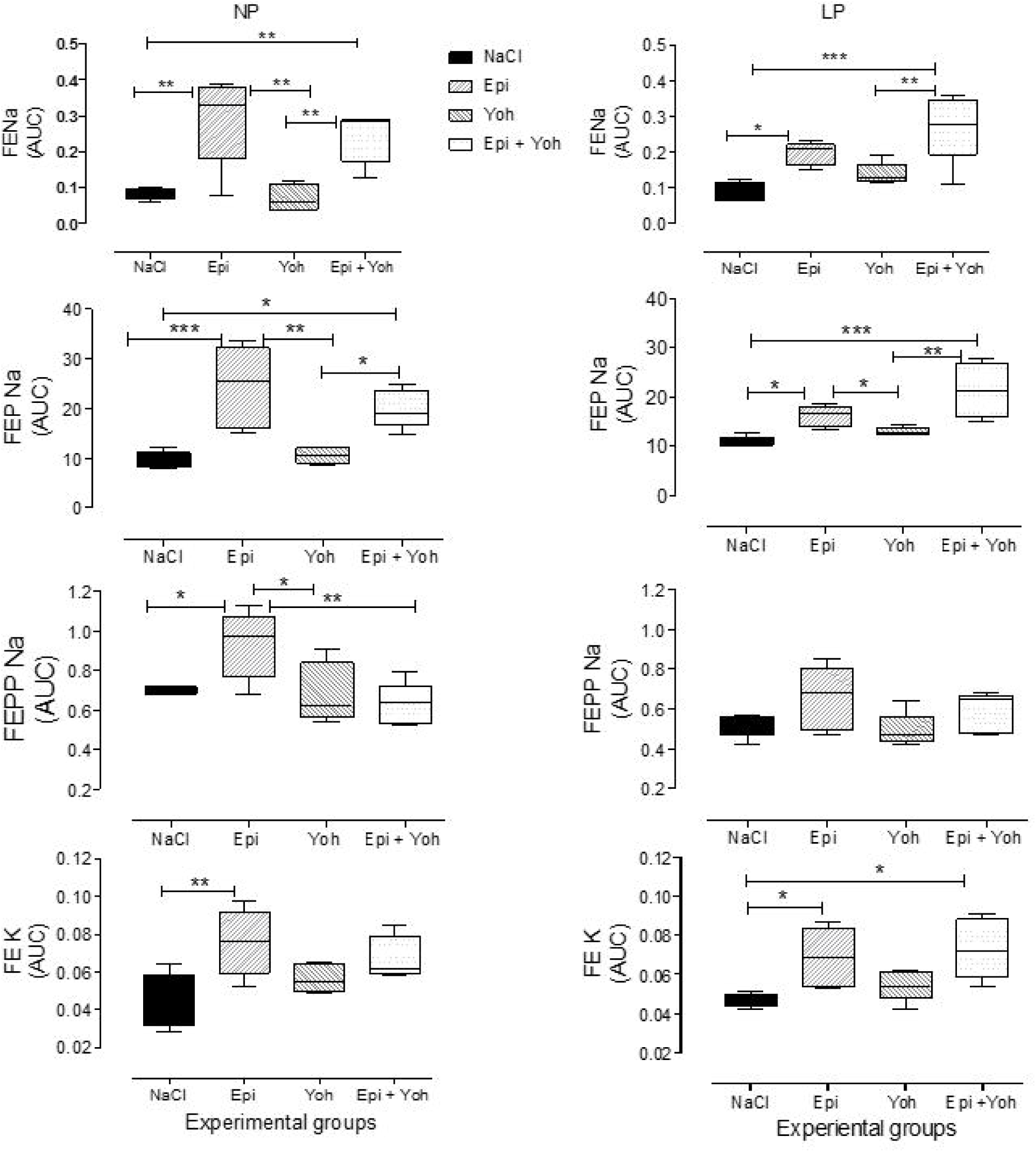
Effect of the lateral ventricle (LV) microinjection of agonist 0.3-µmol epinephrine and/or their α2-adrenoceptor antagonist, 4 nmol yohimbine (Yoh), on fractional sodium excretion (FENa), proximal (FEPNa), post-proximal (FEPPNa) fractional sodium excretion, and fractional potassium excretion (FEK), in NP, compared to age-matched LP offspring. Results are reported as mean ± SD. Data were analyzed using a two-way ANOVA test with post-hoc comparisons by Bonferroni’s contrast test. The level of significance was set at *P < 0.05. The study was controlled by the lateral ventricle (LV) microinjection of 0.15 M NaCl.

## DISCUSSION

The interaction between environmental and genetic factors interfere in ontogenic development, leading to morphofunctional disorders in tissues and organs in adulthood. In this context, gestational protein restriction is followed by low birthweight in rats which in turn, leads to gender-related changes in blood pressure, kidney function, glucose metabolism, and anxiety-like behaviors in male compared to female offspring. The sex hormones contribute to a sexual phenotype dimorphism in the fetal programming model of adult disease by modulating regulatory pathways critical in the long-term control of neural, cardiovascular, and metabolic functions [1-8,35-39]. Thus, this study was conducted only in male rats to ward-off interference from gender differences. The current study confirms a reduced birth weight of rats whose mothers were fed a gestational restricted-protein diet compared to an NP intake [3,4,40]. However, beyond the fourth week of age, body mass in both groups was the same, a phenomenon known as catch-up growth. This effect was associated with a significant enhancement in arterial blood pressure in the LP group. The present investigation also confirmed a pronounced decrease in fractional urinary sodium excretion in maternal protein-restricted offspring (Figure 3). The decreased FENa observed in LP offspring when compared to age-matched NP group, were accompanied by reducing post-proximal tubule sodium rejection, although the creatinine clearance was unchanged and sodium was usually filtered (3,4,41). The decreased renal potassium excretion verified in LP offspring suggest that tubular sodium reabsorption in LP offspring occurs before distal nephron segment.

The precise mechanism underlying the chronic arterial hypertension in offspring induced by maternal low-protein intake has not been identified. Arterial pressure is thought to be controlled by the renal-mediated regulation of fluid and electrolytes. A prior report from our lab has shown higher renal and plasma catecholamine levels in LP offspring when compared to age-matched NP rats. Also, as demonstrated in the previous study [8], the bilateral renal denervation reduced kidney catecholamine concentrations in both NP and LP groups, though the decreased arterial blood pressure was observed only in growth-restricted offspring relative to renal denervated control rats. The enhanced urinary sodium excretion in kidney denervated LP offspring suggests an indirect but close relationship between enhanced renal nerve activity and attenuated sodium excretion in the development of hypertension in LP offspring. These findings are reiterated by a study showing that impaired pelvic neurokinin expression associated with responsiveness of renal sensory receptors in 16-wk old LP offspring are conducive to excess renal reabsorption of sodium and development of hypertension in this programmed model [8]. Otherwise, investigators have demonstrated that administration of adrenergic agonists into different cerebral sites elicits a substantial increase in renal sodium excretion accompanied by decreasing arterial pressure [10,16-18,21]. Thus, we hypothesized that enhanced blood pressure in maternal low-protein intake offspring is associated at least in part, with changes in renal neural activity and reduced urinary sodium excretion that may relate to imbalanced central adrenergic receptors modulation. Here, we evaluated in the concentration-dependent fashion, the effect of cerebro-lateral ventricle administration of adrenergic agonists and/or antagonists, on blood pressure and urinary sodium handling, in 16-wk old LP offspring compared with appropriate age-matched NP controls. Of particular interest, confirming results from different stimulation techniques [10,16-18,21,42-44], this study revealed a rapid, transient, but significant blood pressure decrease after LV microinjection of Epi; this effect was, in turn, attenuated by α1-adrenoceptor antagonist and unchanged by α2-receptor antagonist intracerebroventricular microinjections in NP rats. Additionally, LV epinephrine microinjections, in a dose-dependent fashion, promoted an increase in urinary sodium and potassium excretion over 120 min in NP rats; conversely, the blood pressure and natriuretic response to Epi microinjection into LV were significantly blunted in age-matched LP offspring.

This study confirmed the participation of central nervous system α1 and α2-adrenergic receptors in the regulation of renal sodium and potassium excretion. The increased natriuresis and kaliuresis response to LV Epi microinjection in NP rats were significantly attenuated by previous local injection of prazosin, an α1-adrenergic antagonist. Note that LV microinjection of prazosin inhibited fractional sodium excretion in NP and, to a less extent in LP rats, whereas, the current findings demonstrated that LV pre-injection of yohimbine, an **α2**-adrenergic antagonist, synergically potentiated and normalized the action of intracerebroventricular Epi administration, on renal sodium excretion and blood pressure in age-matched LP offspring. Thus, the study confirmed that epinephrine, when centrally microinjected in conscious rats leads to a very predictable and reproducible natriuretic response accompanied by unchanged glomerular filtration rate, and associated with an increased ion delivery from the proximal tubule incompletely compensated by more distal nephron segments. This effect demonstrates diminished epinephrine graded-fashion responses with a rightward shift of the dose-response curve, providing evidence of down-regulation of target organ responsiveness to LP cerebroventricular stimuli. Despite repeated demonstration of the natriuretic effect of central epinephrine administration to a variety of species, to the best of our knowledge, there has been no previous description of these effects among LP offspring. However, the precise mechanism of this phenomenon remains unclear. Several possibilities might be considered to explain the natriuretic response in this study. First, the central nervous system directly affects renal sodium excretion via neural routes. Second, nephron hemodynamic changes are responsible for alteration of tubule electrolyte handling. Third, the natriuresis results from fluctuations in the level of presumable neural-borne factors which, disrupt sodium and water transporters function in renal tubules.

Moreover, fourth, the attenuated response in LP relative age-matched NP offspring, supposedly be explained by a definite lack of a relationship between centrally adrenergic modulation and/or receptors that blunt the blood pressure and peripheral renal responses. The decreased FENa^+^ responses observed in LP offspring study may result from the interactions of a variety of mechanisms, such as renal arteriolar postglomerular vasoconstriction, renal sympathetic nervous system overexcitability and, by direct tubule transport effects; our previous study has demonstrated increased activity of the Na^+^/K^+^-ATPase pump in the basolateral membrane in LP rats [3,4]. However, in the current study, these effects were not reversed by cerebroventricular adrenergic stimuli.

There is evidence of the importance of renal sympathetic nerve activity in the pathogenesis of experimental models of hypertension [11,28,29]. We have previously demonstrated that the urinary sodium excretion response to central administration of insulin, angiotensin II, hypertonic saline and cholinergic and noradrenergic agonists were strikingly and similarly attenuated in different models of hypertensive rats when compared with age-matched normotensive controls [8,21,35,45,46]. Thus, the significant reduced natriuretic response in LP compared to NP rats may reflect a hyperactive state in the peripheral sympathetic nervous system, including to the kidneys, at least in part, caused by reduced sensory renal activity in LP offspring in adult life [8,29,47,48]. This dysfunctional response could be essential to the development and maintenance of hypertension in maternal protein-restricted offspring.

The adult kidney comprises several filtering units; in some species, total numbers of nephrons are determined before birth [3,31]. In rodents, the permanent metanephric kidney is very immature at birth and, in rats, **about** 20% of the total nephron number is present at birth. There is evidence that any insult, including maternal undernutrition offspring, alters the total number of nephrons and also causes late-onset hypertension [3,4,41]. However, in the present study, it does not seem merely that a reduced nephron number is responsible for the increased blood pressure since we did not observe any significant difference between LP and NP glomerular filtration rate.

It is well known that stimulation of α2-adrenoceptors in the brainstem, in conscious rats, causes a decrease in blood pressure and enhanced urinary sodium excretion selectively mediated by downstream Gαi2, but not Gαi1, Gαi_3_, Gαo, or Gαs, subunit GTP-binding regulatory protein signal transduction pathways [42-44,49,50]. Studies revealed that the brain Gαi2 protein-mediated sympathetic inhibitory renal nerve-dependent path and is of critical importance in the central neural mechanisms activated to maintain fluid and electrolyte homeostasis. The underlying mechanisms by which brain Gαi2-subunit protein-gated pathways induce α2-adrenoreceptor-evoked sodium control in vivo are unclear. Here, given the intimate association between fluid and electrolyte homeostasis and the long-term control of arterial pressure, we may speculate that downregulation of brain Gαi2 protein expression in maternal protein-restricted offspring may lead to high kidney sympathetic drive, renal sodium retention, and the development of renal nerve-dependent hypertension. Our experiments furnished good evidence of the existence of a central adrenergic mechanism consisting of α1 and α2 receptors, which work antagonistically on the regulation of blood pressure and renal sodium excretion. Speculatively, we may suppose that stimulation of central nervous system α2-adrenergic receptors prevents basal increased renal sympathetic overexcitability in conscious LP rats based on two main findings. First, the effect of LV administration of epinephrine on natriuresis is significantly attenuated in gestational protein-restricted offspring. Second, pretreatment with α2-adrenergic receptor antagonists reversed the impact of the LV Epi injection, which demonstrates that central α2-adrenergic receptors are involved in the diminished natriuresis observed for the LP lineage [42-44]. However, we may not discount the possibility that LP neural synapses have more α2-adrenergic receptors than those of NP offspring. It is more likely that natriuresis is a result of reduced renal sympathetic nerve activity and a consequent decrease in renal tubular reabsorption of sodium, and by consequence simultaneously, reduction in the blood pressure in LP offspring. Because adrenergic agonist or antagonists did not alter glomerular filtration rate, changes in glomerular dynamics do not explain the natriuresis. Catecholamines administration into several CNS places of conscious rats increases urinary sodium excretion; the natriuresis is prevented by central α1-adrenergic receptor blockade and potentiated by **α2**-adrenergic receptor blockade [26,27,42-44]. Taking into account the current and previous studies, we may suggest an inhibitory effect of central α2-adrenergic receptors on urinary sodium excretion and an excitatory effect of central α1-adrenoceptors. In conclusion, our results suggest the striking participation of central adrenergic receptors in the renal pathogenesis of elevated blood pressure in LP offspring. Although the precise mechanism of the different natriuretic response of NP and LP rats is still uncertain, these results led us to speculate that inappropriate neural adrenergic pathways might have significant effects on tubule sodium transport, resulting in the inability of the kidneys to control hydrosaline balance, and, consequently, an increase in blood pressure.

*Authors’ contributions:
BVC: Data curation, Investigation, Methodology, Visualization, Writing– original draft; AHC: Data curation, Investigation; PAB: Formal analysis, Methodology, Visualization, Writing–original draft, Writing–review & editing; JAG: Conceptualization, Formal analysis, Funding acquisition, Methodology, Resources, Supervision, Visualization, Writing–original draft, Writing–review & editing

## Acknowledgements

This work was supported by Fundação de Amparo á Pesquisa do Estado de São Paulo (#2013/12486-5), CNPq (#500868/91-3 and 142188/14) and Coordenação de Aperfeiçoamento de Pessoal de Nível Superior (CAPES).

